# Ultrastructure of Precapillary Sphincters and the Neurovascular Unit

**DOI:** 10.1101/2022.12.28.506355

**Authors:** Søren Grubb

## Abstract

Neurons communicate with vasculature to regulate blood flow in the brain. The cells that maintain this are collectively named the neurovascular unit (NVU). This communication, known as neurovascular coupling, is thought to involve astrocytes or molecules that can pass through the astrocytic endfeet. However, the exact mechanism is still unclear. Using large 3D electron microscopy datasets, we can now study the entire NVU in context. In this study, I provide evidence for the role of precapillary sphincters as a hub for neurovascular coupling and endothelial transcytosis, as well as the role of collagen synthesized by fibroblasts in strengthening first-order capillaries. I also show how astrocytic endfeet form a barrier for fluid flow and how the microvasculature of the cortex is not innervated but is surrounded by a surprising organization of parenchymal neuronal processes around penetrating arterioles and arterial-end capillaries in both mouse and human brains.

**Significance statement:** The neurovascular unit (NVU) is made up of various types of cells, including neurons, astrocytes, and endothelial cells, which work together to regulate blood flow in response to changes in neural activity. This process, known as neurovascular coupling, is crucial for ensuring that the brain receives an adequate supply of oxygen and nutrients. This study suggests a novel organization of the NVU and neurovascular coupling. Through ultrastructural analysis, I was able to identify previously unknown relationships between the different types of cells in the NVU. These findings provide new insights into the structure of the NVU and how it functions, which may help researchers develop new strategies for preserving cognitive function and promoting healthy aging.

## Introduction

The Neurovascular Unit (NVU) is a structure that is responsible for the blood-oxygen level dependent (BOLD) response, which links neural activity to changes in blood flow(1). The NVU consists of endothelial cells, vascular mural cells, glial cells, and neurons. While these cell types have been extensively studied(1–5), there is still much to learn about their structures and interactions. Understanding how these cells communicate and what goes wrong during the progression of brain diseases is crucial for developing treatments to halt disease progression.

Neurovascular coupling (NVC) has been an area of interest for over a century. In his Nobel prize lecture, August Krogh asked: “In what way can the capillaries be excited – chemical, electrical, or mechanical? Are they under nervous control, and, if so, by which nerve?”(6).

We have recently described the structure, location and function of brain precapillary sphincters in the brain(7), which are an important part of the NVU as they form a bottleneck for blood flow to the capillary bed and help distribute blood flow between the layers of the cortex. More than half a century ago, Johannes Rhodin studied the ultrastructure of precapillary sphincters in the rabbit thigh-muscle fascia(8), but the ultrastructure of brain precapillary sphincters is still not well understood. His work inspired this study.

In this work, I present novel data on the ultrastructure of the NVU based on a public 3D transmission electron microscopy (EM) dataset visualized by the Neuroglancer website(9), with a focus on brain precapillary sphincters. The 3D EM dataset reveals a distinct organization of neuronal processes around arterial-end microvasculature, the role of astrocytic endfeet in forming a barrier, and the intracellular details of precapillary sphincters.

## Results

This work explored 21 penetrating arterioles (PAs) in a ∼1 mm^3^ part of the mouse visual cortex(9) (tinyurl.com/48vf8pdd) and identified 22 precapillary-sphincters out of 98 studied PA branchpoints, most of them on the first 3 capillary branches of the PAs (Supplementary Table 1, Figure 1a). In the following, I will describe the ultrastructure of the cells that form the NVU, from vascular cells to astrocytes and neurons and compare the ultrastructure with what is currently known about these cell types.

**Figure 1.**
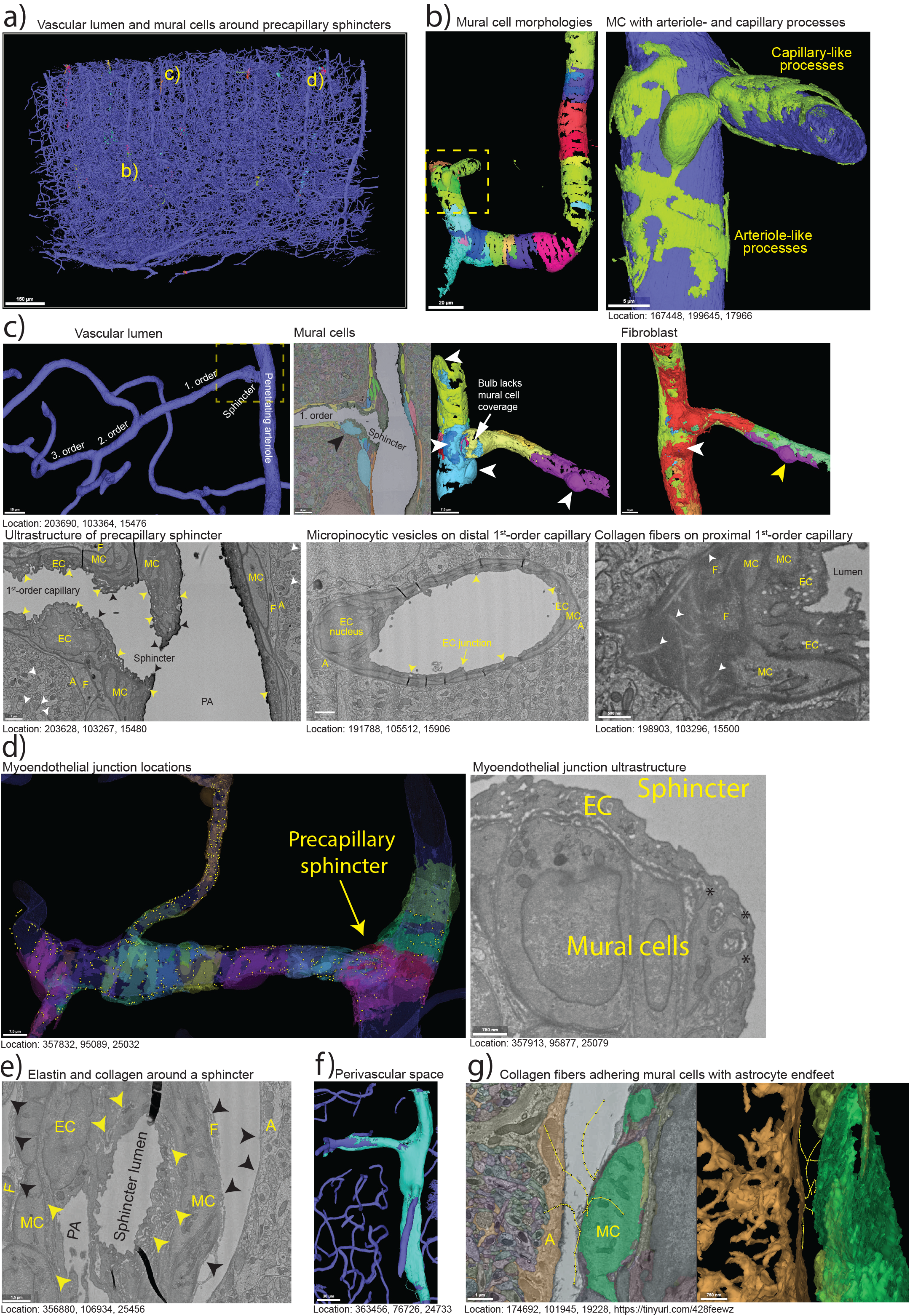
Ultrastructure of the precapillary sphincter. **a)** Overview of the vasculature lumen in the dataset with mural cells around precapillary sphincters visible. The yellow letters indicate where the vessels in Figure 1b, c and d are located. **b)** *Left panel:* Mural cells around a PA are not simple bands but have multiple processes encircling the arteriole. *Right panel:* Mural cells that are situated both on the arteriole and capillary have processes typical of arteriolar mural cells and ensheathing pericytes. **c)** *Upper Left panel:* 3D segmentation of vascular lumen of a PA with 1^st^-3^rd^-order capillaries. The dashed lines indicate the area depicted in the panels to the right. *Upper central panels:* Combined EM and 3D segmentation of mural cells around the PA branchpoint. Black arrowhead points at an endothelial nucleus that lacks coverage by mural cell processes. White arrowheads point at mural cell somas. *Upper right panel:* 3D segmentation of a fibroblast with its soma (white arrowhead) near the branchpoint. The yellow, red, and green segmentation are all a part of the same fibroblast. The fibroblast only partly covers the ensheathing pericyte (yellow arrowhead) on the 1^st^-order capillary. *Lower left panel:* Ultrastructure of the precapillary sphincter showing how layers of mural cell (MC) processes encircle the indentation in contrast to a single layer on the PA and on the 1^st^-order capillary. It also shows how the endothelial cell (EC) nucleus protrudes into the lumen and is in direct contact with the fibroblast (F) or astrocyte (A) endfeet on the abluminal side. The glycocalyx (black arrowheads) is thick and dark on the PA, but lighter and thinner after the precapillary sphincter. The number of EC micropinocytic vesicles is higher immediately downstream of the precapillary sphincter than in the PA. Synapses (white arrowheads) also exist nearby the vasculature. *Lower central panel:* Further down the 1^st^-order capillary the number of micropinocytic vesicles is smaller. See also Video 2. *Lower right panel:* Collagen fibrils (white arrowheads) can be seen in the extracellular matrix abluminal of mural cells at the proximal end of 1^st^-order capillaries. **d)** *Left panel:* 3D segmentation of a PA with a precapillary sphincter and 1^st^-3^rd^-order capillaries. The yellow dots represent individual myoendothelial junctions (peg-and-socket). See also Video 3. *Right panel:* Ultrastructure of the EC and mural cells of the precapillary sphincter showing 3 myoendothelial junctions (*). **e)** Precapillary sphincters have a high amount of elastin (yellow arrowheads) between the endothelial cells (EC) and the mural cells (MC) and collagen fibrils (black arrowheads) between the mural cells and fibroblasts (F). Collagen fibrils are also present in the PVS on the luminal side of astrocyte (A) endfeet. **f)** 3D segmentation of the PVS (turquoise) and the vascular lumen (dark blue). The PVS reaches from the surface of the brain deep into the cortex along PAs but stops at 1^st^-order capillaries. **g)** Tracing of collagen fibrils (yellow lines with dots) crossing the PVS from the surface of mural cells (green) to the astrocyte endfeet (orange).

### Vascular mural cells

The term “mural cell” refers to both vascular smooth-muscle cells and pericytes. Mural cells form the second layer of the blood vessel wall, secondary to endothelial cells. The 3D-ultrastructure of mural cells show a transition of morphologies(2) dependent on the size of the vessels (Figures 1b and 4a). The precapillary sphincter is encircled by contractile mural cells(7), with their soma residing at the arteriole-capillary branchpoint, at the 1^st^-order capillary or at the PA (Figure 1c, Video 1, tinyurl.com/5ce68978) with overlapping processes at the branchpoint. Precapillary sphincters mostly prevail at proximal branches of arterioles in the upper layers of the cortex(7), which is also the case in this dataset (Supplementary Table 1). However, the dataset also contains a precapillary sphincter at a white-matter arteriole (Supp. Fig. 1d).

Intracellular structures of mural cells can also be studied in the MICrONS dataset. The contractile apparatus can be observed in mural cells as faint intracellular lines traversing the cells (Supp. Fig. 1a), like previously shown(8), but becomes increasingly difficult to identify as the capillary branches. At the level of non-contractile thin-strand pericytes, the segmentation is faulty and often combines pericytes with endothelial cells (Supp. Figure 1c). Thin-strand pericytes in the MICrONS dataset were described in the study by Bonney et al.(10), and will, therefore, not be described in detail in this work.

Vesicles and/or caveolae are abundant at the luminal and abluminal plasma membrane of mural cells at PAs and 1^st^-order capillaries. However, like previously reported(11), at higher-order capillaries they are rare and exist mainly on the abluminal side (Supp. Figure 1b).

Elastin is an extracellular-matrix component in the basal lamina of arteries and arterioles that is synthesized by mural cells. It provides elasticity that enables vasculature to dampen pulse-pressure waves traveling from the heart to the microcirculation, known as the “windkessel effect”. Elastin, stained by Alexa633 hydrazide is particularly dense at the precapillary sphincter but terminates there and is not observed in capillaries(7, 12). In the MICrONS dataset, elastin can be observed as a wrinkled light-grey layer with dark borders in pial- and PAs (Figure 1e and Supp Fig. 2a) and at precapillary sphincters (Figure 1e). The elastic fibers consist of fibrillin and elastin. In EM images, fibrillin appears as a dark rim and elastin as a light grey core and is present up until the precapillary sphincter (Supp Fig. 2b), but not beyond that point, consistent with Alexa633 hydrazide staining(7).

These data corroborate what we have found by fluorescence microscopy, that transitional vascular mural cells are a continuum of shapes, and that the precapillary sphincter marks the end of the elastic lamina. Furthermore, they reveal that mural-cell processes overlap at the precapillary sphincter and that a high number of vesicles/caveolae on the luminal side of mural cells are limited to the arterial end.

### Endothelial cells

Endothelial cells form the inner lining of the vasculature in a cobblestone pattern(2). The cells are more elongated in the arterial end than in the venous end. This is not possible to visualize in the MICrONS dataset; however, luminal protrusions at the endothelial-cell junctions are visible as indentations into the vascular lumen segmentation, which allows identification of cell borders despite segmentation problems (Supp. Fig. 1e, tinyurl.com/4hsmkcwk).

As described previously(2, 7), endothelial-cell nuclei are partly uncovered by mural cells immediately downstream of the precapillary sphincter (Figure 1c). Otherwise, most of the arterial-end endothelia is covered.

The luminal side of endothelial cells are coated by a layer of glycoproteins (glycocalyx) which is a part of the tripartite blood-brain barrier(13). The glycocalyx is non-uniform along the vascular tree, with the highest presence at arterioles and with “hot-spots” at arteriole branchpoints(14), perhaps reflecting the amount of shear stress in the tissue. In the MICrONS dataset, the glycocalyx appears darker and/or thicker at arterioles compared with nearby capillaries and venules, with an abrupt transition at the precapillary sphincter (Figure 1c), consistent with a higher density of glycocalyx in arterioles.

Rhodin(8) described how endothelial cells at precapillary sphincters in the rabbit thigh-muscle fascia were ‘rich in pinocytic vesicles, particularly on the luminal side. This is confirmed in the MICrONS dataset, where endothelial cells at precapillary sphincters compared to PAs and downstream capillaries have a high presence of tubular pinocytic vesicles (Figure 1c). They look like what has been found in the hagfish cerebral endothelium(15), and may have a role in fast transcytosis of water and small molecules in a glycocalyx-dependent manner, avoiding lysosomal degradation(16). Ruffles exist in the luminal membrane of these endothelial cells that resemble micropinocytic invaginations and/or macropinocytic membrane protrusions (Figure 1c, Video 2). Rhodin also found that myoendothelial junctions in precapillary sphincters were “more pronounced” and “frequent” compared with arterioles and that roughly 10-20% of the precapillary-sphincter basal endothelial plasma membrane was specialized as myoendothelial junctions(8). Consistent with this, in the MICrONS dataset, myoendothelial junctions are more frequent at precapillary sphincters than at the PAs or 1^st^-order capillaries (Figure 1d, Video 3, tinyurl.com/h6s3pwsh). Interestingly, myoendothelial connections (“peg-and-sockets”) are also abundant at higher-order capillaries where they may have a role in adhering thin strand pericytes to the endothelium(17). Myoendothelial junctions are enriched along the edges of mural-cell processes. Contrary to Ornelas et al., who looked at “pegs-and-sockets” near an ascending venule(17), in the arterial end they are not particularly enriched near the mural-cell somas.

These data suggest that precapillary sphincters are a hotspot for myoendothelial communication, changes in glycocalyx density, and fast transcytosis of water and small molecules from the blood across the endothelium.

### Perivascular fibroblasts

Perivascular fibroblasts are distinct from pericytes and may be a continuation of pial fibroblasts(18). They can be identified by their long slender processes and their location, which is abluminal to mural cells and luminal to astrocyte endfeet and/or macrophages. Contrary to macrophages they have only few lysosomes and phagosomes, and I have not yet found a fibroblast that did not have a primary cilium.

Collagen type I fibrils and fibers are synthesized and secreted by fibroblasts(19) to scaffold and strengthen the vasculature to withstand blood pressure. These fibrils/fibers exist on the abluminal side of contractile mural cells(7) and are visible in the MICrONS dataset as thin parallel fibrils with a diameter of approximately 50-100 nm with cross-striations (Figure 1e and g, Supp. Fig. 2a and c, Video 4). In the MICrONS dataset, invaginating pockets (fibripositors) with terminal collagen fibrils are found on the luminal side of fibroblasts (Supp. Fig. 2d), indicating collagen fibril assembly (Supp. Fig. 2d and Video 4).

An example of a perivascular fibroblast next to a PA-branchpoint mural cell has been observed previously in the MICrONS dataset(10); however, the cytoarchitecture of that cell included dark-grey lysosomes and phagosomes, suggesting that it is likely a macrophage and not a fibroblast. Conversely, a fibroblast is found on the luminal side of that macrophage (Supp. Fig. 2e, tinyurl.com/4s6zja7z). Perivascular-fibroblast somas are found on pial- and PAs, ascending venules and on 1^st^-order capillaries but rarely beyond that point (Figures 1c and 4a), with processes reaching 3^rd^-order capillaries. Where the fibroblast processes exist, collagen fibrils can also be found (Figure 1c).

A fibroblast- and/or macrophage-soma location next to the precapillary-sphincter mural cell is often associated with an indentation in the PA lumen (Supp. Fig. 2f, tinyurl.com/dhwf8jtt), this has also been observed *in vivo*(7).

Perivascular spaces (PVS) can be found around PAs and are sometimes visible in the segmentation (Figure 1f). Collagen fibrils are also present at the luminal side of the astrocytic endfeet, which is visible where there is a PVS (Figure 1g, tinyurl.com/43f92u7m). Some collagen fibrils even connect the abluminal mural-cell extracellular matrix with the parenchymal basal lamina. Only in places devoid of mural-cell coverage are fibroblasts in direct contact with the endothelial basal lamina (Supp. Fig. 2g).

These data show how fibroblasts synthesize collagen fibrils to strengthen the PAs and 1^st^-order capillaries and how the fibrils reach across the PVS to connect mural cells with astrocytic endfeet.

### Astrocytes

Astrocytes are regularly illustrated as a star-shaped cell body located in the parenchyma with long processes that form endfeet on blood vessels. In a recent review, we argued that this may not always be the case, as astrocytes may have their soma situated directly on blood vessels(2). In the MICrONS dataset, it appears that for large vessels (both arterioles and venules), this is more a rule than an exception (Figure 2a, 4b and Video 5), whereas, for capillaries, the astrocyte soma is often located in the parenchyma (Figure 2a). From the astrocyte soma, large processes stretch out in the parenchyma and divide into smaller processes that come in close contact with neuronal synapses.

**Figure 2.**
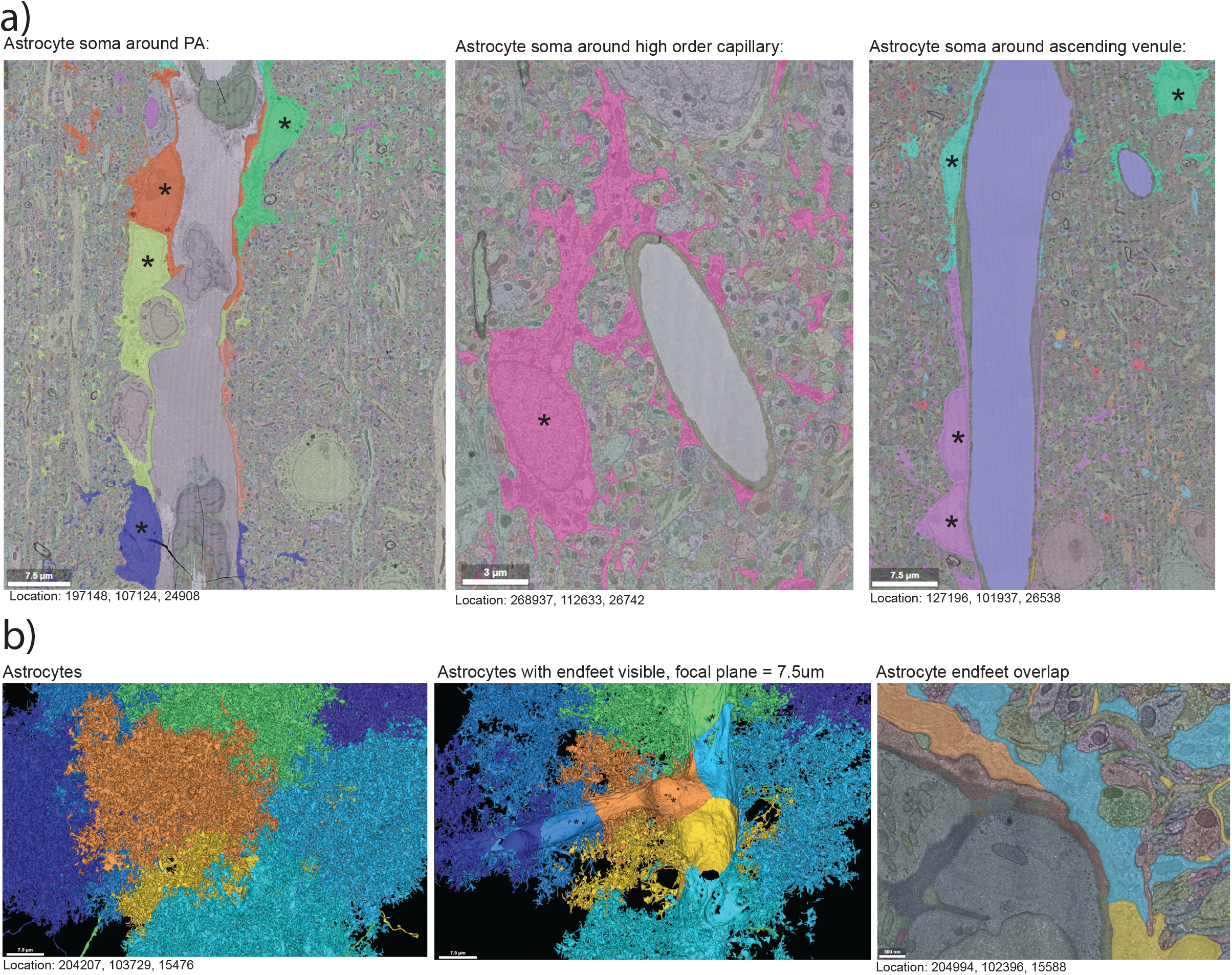
Astrocyte ultrastructure. **a)** Ultrastructure shows that astrocyte somas (*) frequently forms the “endfeet” on PAs (*left panel*) and ascending venules (*right panel*) but the usual depiction of an astrocyte in the parenchyma with endfeet encircling the vessel is typically found on capillaries (*central panel*). **b)** *Left panel:* 3D segmentation of 8 astrocytes around a PA branchpoint shows that the astrocytes each cover a discrete parenchymal volume. *Central panel:* By reducing the focal plane to 7.5 µm we can see the endfoot coverage is also discrete with sharp lines where the endfeet meet. Small holes in the center of the endfeet are segmentation errors, not perforations. *Right panel:* Ultrastructure shows a large overlap between astrocyte endfeet, without any visible gaps.

Astrocytes have well-defined domains that are populated by the fine processes and endfeet (Figure 2b, tinyurl.com/7spjzjz5). The astrocyte endfeet overlap (Figure 2b) and add to the barrier function of the blood-brain barrier by restricting diffusion(20). Occasionally, one may find openings between astrocytic endfeet where a single axonal bouton protrudes to contact a blood vessel, but I have only observed that in high-order capillaries (Examples: tinyurl.com/47rnmmfs and tinyurl.com/37vry6nu). Notably, at some endfeet the astrocyte membrane becomes extremely thin to give the impression of incomplete coverage (Figure 2b). However, close inspection of the EM data reveals that the features originate from segmentation artifacts caused by the collapsed endfoot plasma membrane blending in with the extracellular matrix of similar electron density.

Together, these examples show how, contrary to what is generally thought, astrocyte soma form a part of the endfeet at PAs and ascending venules and that astrocytic endfeet form a tightly overlapping barrier which may limit paracellular communication.

### Precapillary sphincter innervation

Perivascular nerve fibers originating from the sympathetic nervous system innervate cerebrovascular arteries (21), and it is thought that they exist on vessels up until their entry into the brain parenchyma. Rhodin found “abundant” unmyelinated nerve endings near the precapillary sphincter in the rabbit thigh fascia with axonal-terminal boutons containing vesicles with granules(8), indicating direct neuronal modulation of precapillary-sphincter function. However, in the MICrONS dataset, I found no perivascular innervation of cortical PAs, precapillary sphincters or even of the largest arterioles (Supp. Fig. 3a). All neuronal processes reside on the parenchymal side of astrocytic endfeet, arguing against direct nerve control of vascular function in the cortex.

Interestingly, small unmyelinated axonal processes abound closest to the astrocyte endfeet of PAs and 1^st^-order capillaries with synaptic vesicle-filled boutons facing the endfeet, while dendrites and dendritic spines most often lie in the second row (Supp. Fig. 3b, Video 2, Axons: tinyurl.com/er6yumyn, Dendrites: tinyurl.com/4db9pyfn). At high-order capillaries there are relatively more dendrites closest to the vasculature. Unfortunately, segmentations of many of the axonal processes are not complete in the MICrONS dataset, and it is therefore difficult to characterize what type of neurons the processes originate from. However, of the neuronal processes near the 1^st^-order capillaries where the origin is possible to trace, one can find: spiny dendrites of pyramidal cells as well as processes and somas of many types of interneurons, including Martinotti cells, chandelier cells and oligodendrocyte precursor cells (Supp. Fig. 3c). These data show how contrary to peripheral microvasculature, brain precapillary sphincters are not innervated by perivascular nerve fibers. All neuronal processes reside at the parenchymal side of astrocytic endfeet.

### Neuronal processes are aligned perpendicular to the vascular direction

While investigating the NVU ultrastructure in the MICrONS dataset, it became clear that neuronal processes closest to astrocyte endfeet at PAs and 1^st^-order capillaries are surprisingly organized. Neuronal processes are arranged perpendicular to the vessel direction and curb around the vessel, especially near the precapillary sphincter (Figures 3a and 4b and Video 6). It seems that this arrangement is independent of parenchymal-orientation or depth of the vessel, as it can be found both on PAs and capillaries that branch in different angles (Figure 3a) both shallow- and deep in the cortex (Figure 3b). At capillary bifurcations, neuronal processes appear more chaotic, but the organization is re-established on daughter branches (Figure 3a).

**Figure 3.**
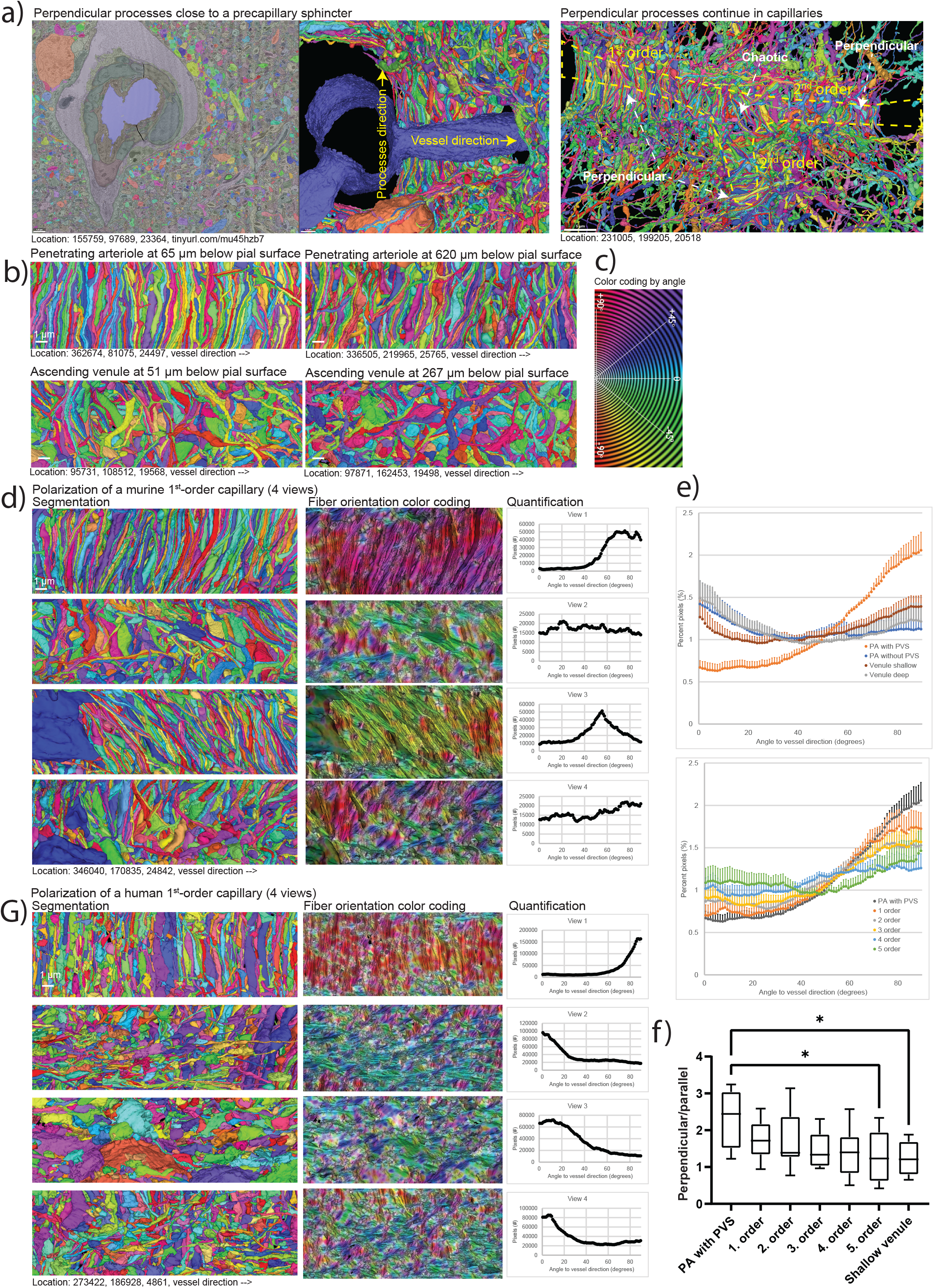
Neuronal processes around the brain microvasculature. **a)** *Left panel:* Ultrastructure of a PA branchpoint (lumen is blue) where the neuronal processes closest to the astrocyte endfeet have been made visible in the 3D segmentation (right half). It shows that neuronal processes are aligned at a 90° angle to the vessel direction. *Right panel:* 3D segmentation of neuronal processes around both 1^st^- and 2^nd^-order capillaries show that as the vessel bifurcates (dashed yellow line), the perpendicular arrangement is disrupted, but is reestablished after the bifurcation. **b)** The perpendicular arrangement is present in PAs, but not in ascending venules. *Upper panels:* Example of the arrangement of neuronal processes around a PA at 65 and 620 µm below the brain surface. *Lower panels:* Example of the arrangement of neuronal processes around an ascending venule at 51 and 267 µm below the brain surface. In this example, the neuronal processes are only shown from one viewing angle. Scalebar = 1 µm. **c)** Indication of angular color-coding of the ImageJ plugin “OrientationJ” that was used to quantify the perpendicular arrangement of neuronal processes. **d)** Example of 4 views of the closest neuronal processes to the astrocyte endfeet of a 1^st^-order capillary. *Left panels:* 3D segmentation of the neuronal processes, the vessel is oriented horizontally. *Central panels:* Color coding of angles of neuronal processes to the vessel orientation. *Right panels:* Quantification of the pixel distribution of angels in the 4 views. **e)** *Upper panel:* Mean percent of pixels of all 4 views at different angles to vessel direction for PAs with or without PVS (PVS) and for ascending venules shallow or deep in the cortex. Error bars are SEM, n = 6-7. *Lower panel:* Mean percent of pixels at different angles to vessel direction for PAs with PVS and 1^st^-5^th^-order capillaries. Error bars are SEM, n = 7-8. **f)** Box and whisker plot of ratio between perpendicular (>45°) and aligned (<45°) processes. Tested with one-way ANOVA, n = 6-7. Two outliers were excluded: 1. order = 4.99 and 3. order = 4.12. **g)** Same as d) but for a 1^st^-order capillary from a human brain sample.

**Figure 4.**
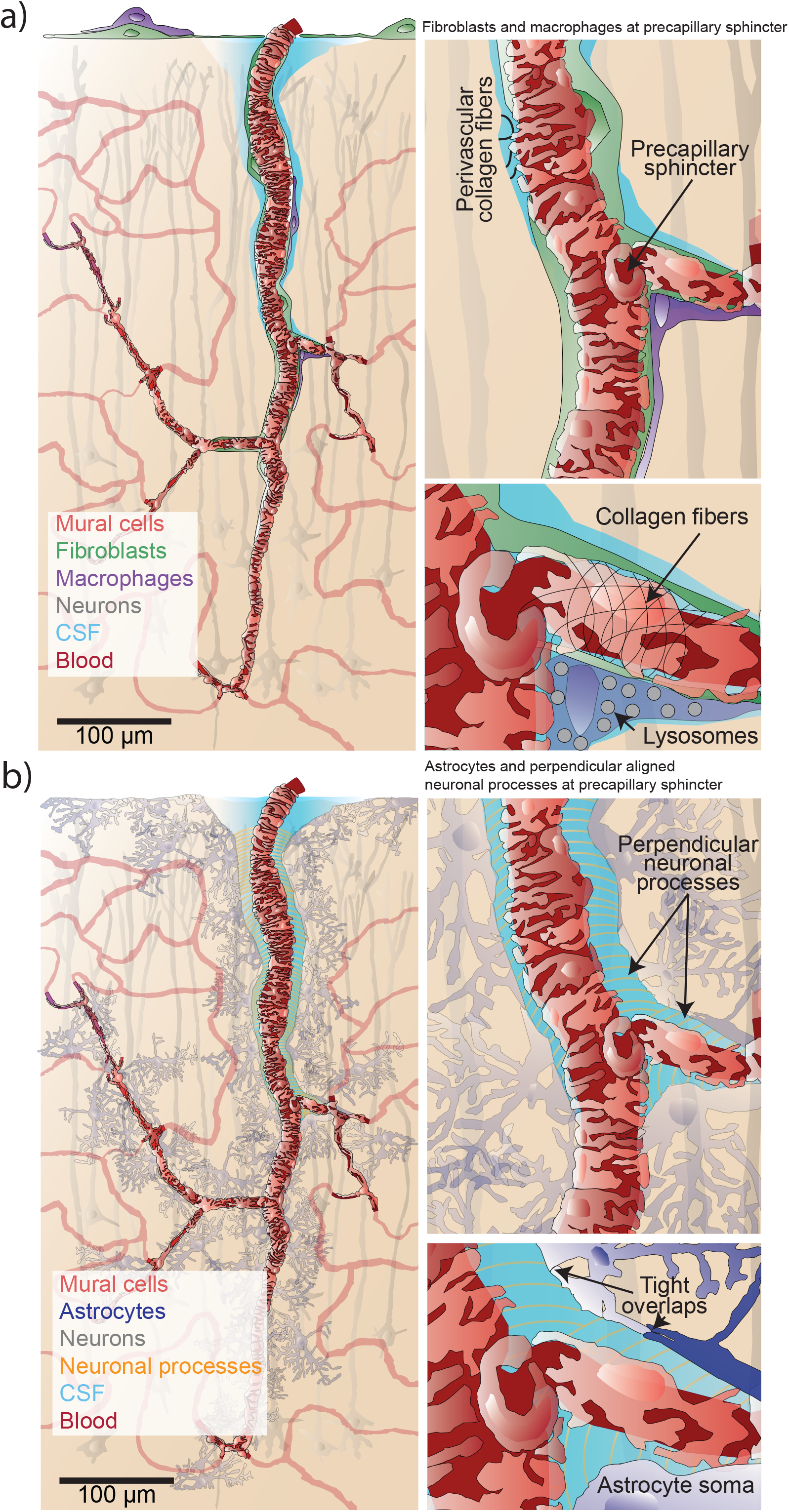
Illustrations of the neurovascular unit. **a)** *Left panel:* Illustration of a PA that branch to a precapillary sphincter. The cell types included are mural cells, neurons, perivascular-fibroblasts and macrophages, pial fibroblasts and a meningeal macrophage. *Right panel:* Enlargement of the precapillary sphincter branchpoint shows how fibroblast- and macrophage processes reach to the 1^st^ order capillary. **b)** *Left panel:* Shows how astrocyte endfeet cover the entire perivacular space and glia limitans. Notice how astrocyte cell bodies are a part of the endfeet on the PA. *Right panel:* Enlargement of the precapillary sphincter branchpoint shows how perpendicular aligned neuronal processes exist both on the PA where there is a PVS and on arterial-end capillaries that also may have a PVS.

It seems this perpendicular organization is limited to one half of the perivascular wall, whereas the second half often has one or more cell bodies taking up space (or thicker astrocytic endfeet) (Figure 3d). These can be either astrocyte cell bodies (Supp. Fig. 4a, tinyurl.com/mr3ry962), microglia (Supp. Fig. 4a, tinyurl.com/2p8288sn), neuronal cell bodies (Supp. Fig. 4a, tinyurl.com/2p8bwynj) or oligodendrocyte precursor cells (Supp. Fig. 4a, tinyurl.com/bdh5fwx6).

Interestingly, the cell bodies taking up space are often abnormally elongated (Supp. Fig. 4a). Occasionally, in the space between the two halves of the perivascular wall, neuronal processes can be seen following the vessel direction (Figure 3d and g, view 2 and 4).

Synapses are present throughout the parenchyma and can therefore also be found near the arterioles and 1^st^-order capillaries (Figure 1c); however, axonal boutons seem more concentrated on the second half of the perivascular wall further away from the vasculature (Figure 3d and g, view 3 and Video 6).

To quantify the arrangement of neuronal processes closest to the astrocyte endfeet, their orientation was analyzed by averaging 4 different views of each segment. This was done for PAs and 1^st^ – 5^th^-order capillaries, as well as ascending venules. If the processes follow the vessel direction, their angles approximate 0° and if they are perpendicular, their angles are close to 90°. Most of the capillary trees analyzed have around 8 capillary bifurcations between the PA and ascending venule as the shortest distance. I found no capillary trees connecting PAs with ascending venules by less than 5 bifurcations. This is the reason for including 5 capillary orders in the analysis.

Neuronal processes in the perivascular wall of PAs with PVSs had a significantly higher incidence of processes going perpendicular to the vessel direction (Figure 3b and e). Interestingly, PAs without PVSs, which are most often found deeper in the cortex and are most often also smaller in lumen diameter, did not have neuronal processes perpendicular to the vessel direction (Figure 3b). In one case, a large PA had a PVS reaching 489 µm and maintained a large diameter to this depth due to lack of bifurcations; it, too had neuronal processes arranged perpendicular to vessel direction, even deep in the cortex (Video 6, tinyurl.com/4fbfmh5s). This argues against the perpendicular arrangement being merely caused by horizontally oriented neuronal processes in the molecular layer of the cortex. Corroborating this, venules found either near the surface or deep in the cortex do not generally have neuronal processes perpendicular with the vessel direction (Figure 3b and e) and venules do not have any PVSs, except for the first few micrometers below the pial surface.

The perpendicular arrangement decreases with capillary bifurcations (Figure 3g and f), and at 5^th^-order capillaries the processes are significantly less rectified than at PAs with PVS.

These remarkable data show how neuronal processes closest to the astrocytic endfeet are arranged perpendicular to vessel direction at PAs with PVS but also on arterial-end capillaries.

### The perpendicular arrangement is translational to human cortical vasculature

In the “H01-release” human cortex dataset(22), also available via Neuroglancer, the same arrangement of neuronal processes half-circling the parenchymal-perivascular wall exists (Figure 3g and Video 7). This suggests the arrangement is not specific to mice or an artifact of the MICrONS sample.

The segmentation in the H01-release dataset is much more fragmented and inaccurate, and astrocytic endfeet are severely swollen, making it difficult to recognize vascular cells(10). However, based on the presence of contractile elements in the mural cells (Supp. Fig. 4b), a PA with a 1^st^-order-capillary branchpoint could be identified. The 3D segmentation shows perpendicular processes on one side of the 1^st^-order capillary, and on the opposite side 6 oligodendrocyte cell bodies sit back-to-back (Supp. Fig. 4c, tinyurl.com/2yc3znte). In between the two sides, neuronal processes align with the vessel direction (Figure 3g, Video 7).

Segmentation of a human ascending venule did not show any alignment of neuronal processes (Supp. Fig. 4d, Video 7), confirming that the polarization and perpendicular arrangement of neuronal processes is specific for the arteriolar side and is preserved between mice and humans.

These data show that the same arrangement of neuronal processes perpendicular to vessel direction exists in the arterial end of human cortical vasculature and may therefore be translatable from mouse to human.

## Discussion

### Transitional mural cells are a continuum of shapes

Mural cells provide blood-vessels with a tonus, which is the prerequisite for blood-flow regulation. I have shown here and previously with coworkers(2, 7) that mural cells on PAs and 1^st^-order capillaries are a continuum of shapes (Figure 1a-c and 4a). It is therefore an oversimplification to categorize PA mural cells as smooth-muscle cells. Furthermore, it is inconsistent with the original definition of pericytes by Zimmermann, that all transitional forms from spindle-shaped smooth-muscle cells (not included) to capillary thin-strand pericytes should be named pericytes(2, 23). Mural cells up until 4^th^-order capillaries may be contractile, but the definition of capillaries/precapillaries/arterioles in the transitional segment is also debated(2).

In the MICrONS dataset, contractile elements may be observed in mural cells of arterioles and 1^st^ – 4^th^-order capillaries (Supp. Fig. 1a) but not beyond. Contractility has been suggested for pericytes beyond the 4^th^-order capillary(24, 25) (thin strand pericytes), which is a very slow process that is thought to involve cytoskeletal elements like g- or f-actin. However, this theory is mostly based on dubious α-SMA labeling in a highly referenced study(26) and confusing differences in capillary-order nomenclature between retina and brain(27).

### The ultrastructure of precapillary sphincters

Precapillary sphincters have been an overlooked feature of the cerebral microvasculature(28), until we described their presence and function by *in vivo* two-photon imaging and immunohistochemistry(2, 7). The MICrONS dataset provides an opportunity to describe the ultrastructure of precapillary sphincters and the NVU in context of cortical depth and microvascular hierarchy.

An example of a PA branchpoint with mural cells encircling it has been shown previously in the MICrONS dataset(10) (Supp. Fig. 2e). However, though the branchpoint does have a mural-cell soma with encircling processes it does not fit our definition(7) of a precapillary sphincter due to the lack of lumen indentation at the branchpoint (Supplementary Table 1, arteriole 2 branch 1).

The mural cells encircling the precapillary sphincters and the overlapping mural-cell processes at the branchpoint indentation are consistent with a high contractility of precapillary sphincters(7, 29), and supports the notion that precapillary sphincters are central to blood-flow regulation. In the following, the main findings from the ultrastructure of precapillary sphincters will be described.

### The precapillary sphincter may be important for transcytosis and fluid uptake

The precapillary sphincter is a place of large changes in blood- and hydrostatic pressure(7), suggesting it is a hotspot for fluid filtration from the blood to the PVS, which in most tissues happens paracellularly. However, in the brain, paracellular transport is severely restricted due to endothelial tight junctions; therefore, transcytosis may have a role in fluid uptake from the blood to the PVS. Transcytosis is normally suppressed in brain-endothelial cells, except for in disease states(30), and a high number of vesicles may indicate damage to the cells, e.g., during the fixation process. However, because the presence of tubular micropinocytic vesicles is mostly confined to the precapillary-sphincter endothelium(8) (Figure 1c); and because there are generally no other indications of cellular damage, like swollen mitochondria or swollen astrocytic endfeet, it suggests that the high amount of tubular micropinocytic vesicles in precapillary-sphincter endothelial cells are related to the location rather than an artifact. Corroborating this, tubular vesicles in brain endothelial cells are hypothesized to be involved in fast transcytosis facilitated by PACSIN-2(16). Furthermore, a sudden change in glycocalyx composition (Figure 1c) may influence transcytosis(16). Finally, the luminal-membrane invaginations of 1^st^-order capillary endothelial cells (Figure 1c, Video 2) suggest the transcytosis is directed from the blood towards the brain, as is required for fluid uptake.

### Myoendothelial junctions are more abundant at the precapillary sphincter

Rhodin speculated that myoendothelial junctions are more than merely a structural component that serves to stabilize the microvascular wall(8). Indeed, myoendothelial junctions are places where contractile mural cells connect with endothelial cells via gap junctions(31). The hyperpolarizing signal in NVC that is conveyed by inward rectifier K^+^ channels in the endothelial cells may spread via myoendothelial gap junctions to contractile mural cells to elicit a vasodilation(32–34). The abundance of myoendothelial junctions reaching across the elastic lamina at the precapillary sphincter (Figure 1d) could explain why proximal 1^st^-order capillaries react earliest to nerve activity(35).

Myoendothelial junctions exist mostly along the edges of the mural-cell processes and can be either endothelial- or mural-cell protrusions or both. Whether all these myoendothelial junctions contain gap junctions and which connexins they contain is uncertain(36). Some of them may simply be “peg-and-socket” attachment points(17). However, especially 1^st^-order capillaries are locations where myoendothelial coupling is expected to be strong(35, 37) and the abundancy of myoendothelial junctions at precapillary sphincters indicate that they are important locations for NVC.

### The elastic lamina ends at the precapillary sphincter

The elastin ultrastructure can be observed in arterioles in the MICrONS dataset, which confirms the termination of elastic lamina at the precapillary sphincter (Figure 1e and Supp. Fig. 2b), as previously reported(7). In contrast, Rhodin concluded that there are no elastic components present in relation to the precapillary-sphincter ultrastructure in the rabbit-thigh-muscle fascia(8), suggesting a different need for microvascular elasticity compared with the brain. This could be ascribed to differences in the need to dampen pulse-pressure waves in the two tissues, as thigh-muscle fascia is less perfused and more distal to the heart compared with the brain.

Interestingly, an explanation for functional K_ir_2.1 in arterioles but not capillaries of Cerebral Autosomal Dominant Arteriopathy with Sub-cortical Infarcts and Leukoencephalopathy (CADASIL) mice(38), could be the presence of elastin in arterioles but not in capillaries, as TIMP3, an inhibitor of K_ir_2.1 channel activity which is accumulated in CADASIL mice, binds with high affinity to elastin, preventing the endothelium from being exposed TIMP3(38, 39). It would be interesting to see whether differences in brain- and peripheral microvascular elastin expression affects the outcomes of CADASIL.

### Perivascular collagen fibrils may couple arteriole constriction with astrocyte activity

While elastin coverage terminates at the precapillary sphincter, fibroblast processes enwrap the 1^st^-order capillaries and collagen fibrils may also be found there (Figure 1c, Video 2) supporting the capillary structurally. These fibrils may also limit the maximal dilation of capillaries(7). On PAs, collagen fibrils that reach astrocytic-endfeet basement membrane across the PVS (Figure 1g) may have a role in triggering mechanosensitive membrane proteins (like TRPV4 channels) and thereby increasing Ca^2+^ in astrocytic endfeet, which releases vasodilators as stretch-mediated feedback, causing oscillations of arteriole diameter(40). Conversely, a recent study has suggested that macrophages degrade these PVS-crossing collagen fibrils and shown that depletion of macrophages limit arterial motion(41).

### Neurovascular coupling is astrocyte dependent

NVC consists of several pathways, like nitric oxide-, glutamate- and purinergic-signaling pathways (2, 41). These can be divided into gasotransmitters (e.g., NO, CO, and H_2_S), neurotransmitters (e.g., glutamate and acetycholine) and neuromodulators (e.g., ATP) and involve intracellular-Ca^2+^ signaling and electrical signaling by changes in membrane ion conductance or electrochemical gradients.

As mentioned above, it is questionable whether NVC happens paracellularly in between the tightly overlapping astrocytic endfeet (Figure 2b), suggesting that neuronal processes signal either via astrocytic endfeet(4, 5) or via gasotransmitters that can pass through the endfeet unhindered.

NO is hydrophobic and highly diffusible and is thought to be the main mediator of vasodilation(42); however, in the cerebral cortex, NO is a modulator (a permissive role) rather than a mediator of NVC(43). Pathways involving astrocytic-endfoot signaling are the strongest candidates for mediators of NVC, at least for capillaries(4).

Astrocyte-dependent pathways involve intracellular Ca^2+^ signaling, which can lead to either a vascular dilation or constriction depending on the level of NO and brain metabolic elements(44). However, only a third of the NVC can be accounted for by known mechanisms, suggesting the involvement of a yet unidentified mechanism(42).

### Perpendicularly aligned neuronal processes may enhance neurovascular coupling

The finding that parenchymal nerve processes are aligned perpendicular to blood vessel direction in the arterial end where there is a PVS (Figure 3), but also on the contractile 1^st^-4^th^-order capillaries where there is generally no PVS, raises the question whether this arrangement has a role in enhancing NVC?

Perpendicularly aligned neuronal processes could possibly **I)** allow more processes to be near the vessel; **II)** minimize the extracellular space between them (compared with a more chaotic arrangement) and thereby locally concentrate the extracellular K^+^ released by action potentials leading to increased K^+^ siphoning by astrocytes (although, astrocytic K^+^ siphoning is not likely a major mechanism of NVC(45)); **III)** minimize the distance that membrane-permeable molecules like NO, O_2_ or CO_2_ need to diffuse to reach their targets; **IV**) minimize the distance of signaling through the astrocyte endfeet; **V)** better sense and react to blood-flow changes (the haemoneural hypothesis(3)). Furthermore, the mentioned polarization suggests that synapses (and therefore neurotransmitter release) are concentrated in certain areas near the vessel.

While these suggestions could explain how the alignment of neuronal processes would be beneficial, it does not explain why the perpendicular alignment is mostly found around PAs with a PVS. What causes these perpendicular alignments is also not clear, it could be caused by vascular wall signaling during migration of the neuronal processes, be controlled by astrocyte processes or be influenced by perivascular pumping. Alternatively, non-aligned processes could be pruned by microglia. Whatever the cause and mechanism is (if any), much work is needed to clarify this.

To summarize, the ultrastructure of the NVU reveals that **I)** the precapillary sphincter is central to the regulation of blood flow and likely also water uptake; **II)** the barrier function of astrocytic endfeet has likely been underestimated; **III)** perpendicularly aligned neuronal processes encircle PAs and 1-4^th^ order capillaries which may enhance NVC.

## Conclusion

Capillaries may be stimulated in several ways, including chemical signaling pathways involving neurotransmitters and gasotransmitters, electrical signals caused by changes in ion conductance and electrochemical gradients across the plasma membrane, and mechanical signals generated by the stretching of vascular mechanosensitive ion channels and receptors. Central to this communication are the precapillary sphincters and 1^st^-order capillaries. In the peripheral microvasculature, capillaries may be directly controlled by the nervous system, but in the cortex, they are indirectly controlled by signaling through astrocytic endfeet or diffusible ions and molecules which may be optimized by a perpendicular arrangement of neuronal processes around the vasculature.

## Methods

### Public volume electron microscopy datasets

The primary dataset used in this work is the MICrONS dataset provided by the Allen institute(9) and composed of an approximately 1 mm^3^ chunk of the mouse (male P75-87) visual cortex, including parts of the primary visual cortex and higher visual areas for all cortical layers except extremes of L1. The dimensions are (*in vivo*) 1.3 mm mediolateral, 0.87 mm anterior-posterior and 0.82 mm radial. The resolution was ∼4 nm but downsampled to 8 nm/pixel, and the slice thickness was 40 nm. This dataset was also used in the study by Bonney et al.(10), where they annotated the PAs and ascending venules. However, the section consists of two subvolumes, one that contains 65% of the total volume and another that contains 35% of the total volume. Here is a vascular annotation for both volumes: tinyurl.com/48vf8pdd. The MICrONS consortium has used the Google-developed Neuroglancer web-based application to visualize the segmented data. A guide to navigating the tool can be found on the MICrONS website: microns-explorer.org/visualization. Each segment has a unique code consisting of 18 numbers, and these can be copied out of the web application to save your work or pasted back into Neuroglancer. The activated segments and viewer settings are included in the URL, so the URL can become extremely long. Therefore, URL’s have been shortened using tinyurl.com. Furthermore, location data have been provided for each figure, it is read as XYZ coordinates and can be pasted into the top left corner of the Neuroglancer website. Movies were created by taking multiple screenshots, using a macro made in iCue – a gamer keyboard software. The screenshots were cropped, stacked, and converted to “.avi” using ImageJ. To test whether some findings were present in a human dataset, the public H01-release dataset(22) was used, which is also available to visualize in Neuroglancer. The dataset consists of a temporal lobe cortical fragment from a 45-year-old patient with drug-resistant epilepsy. The cortical fragment was not abnormal, but the underlying hippocampus was sclerotic(22). The dataset consists of 5000 slices of 30 nm thickness in an area of 4 mm^2^ spanning from the pial surface to the white matter. As mentioned by Bonney et al.(10), the vascular lumen is collapsed, and the astrocyte endfeet are swollen, making it difficult to characterize vascular cells.

### ImageJ and OrientationJ

For analysis of the orientation of neuronal processes, an ImageJ plugin OrientationJ was used on screenshots taken from 4 different views of the parenchymal-perivascular wall (excluding astrocytes). The screenshots were acquired by first right-clicking on a neuronal process in the center of the segmentation to move to that spot, and then clicking “z” to align the orientation to one of the axes. The “left/right” hotkeys and “e” or “r” hotkeys were used to align the blood vessel with the plane of focus and to make it horizontal. The “alt+scroll” hotkey was used to change the focal plane to make the neuronal processes visible and then a screenshot was made. This protocol was repeated for the following three views, each perpendicular to the next. The screenshots were cropped to the diameter of the parenchymal-perivascular wall (excluding astrocytes), and the length of the segmentation and changed to 8 bits using ImageJ. With the settings used, the screenshots had a resolution of 91-97 pixels per micrometer. The crop sizes varied depending on the diameter between the segmented neuronal processes and the length of the activated segmentations. “OrientationJ – Analysis” was used to visualize the orientation of neuronal fibers in the perivascular wall, and “OrientationJ – Distribution” was used to quantify it. For both, a local window of 10 pixels and a Gaussian gradient was used. Positive angles (0.5° to 89.5°) and negative angles (−0.5° to -89.5°) angles were averaged, to find the angles different from the vessel direction (0°).

### Limitations

Although the amount of swelling is limited, the MICrONS dataset does contain some swollen astrocytes. In contrast the human dataset has severely swollen astrocytic endfeet, which makes the identification of arterioles and capillary orders difficult and the characterization of the NVU problematic. Furthermore, chemical fixation will reduce the extracellular space and possibly also the PVS.

## Supporting information

Movie 1 - Mural cell morphology

Movie 2 - Serial EM of Precapillary Sphincter

Movie 3 - Myoendothelial Junction annotation

Movie 4 - Serial EM of Fibroblast

Movie 5 - Astrocyte Cell Bodies around Arteriole

Movie 6 - Perpendicularly aligned Neuronal Processes (mouse)

Movie 7 - - Perpendicularly aligned Neuronal Processes (human)

## Acknowledgements

I would like to acknowledge Professor Martin Lauritzen for letting me work on an independent project, and for helpful comments. Thanks to the Novo Nordisk foundation for supporting me indirectly via Professor Martin Lauritzen.

## Supplementary figure texts

**Supplementary Figure 1**

**a)** *Left panel:* Ultrastructure of an arteriolar contractile mural cell showing contractile elements and abluminal coated vesicles. *Right panel:* Mural cell on a 4^th^-order capillary showing a few lines that could be contractile elements. **b)** *Left and right panels:* Vesicles and caveolae can be found near the luminal and abluminal membrane of mural cells on precapillary sphincters and PAs. *Central panel:* In capillaries vesicles and caveolae are mainly found on the abluminal membrane. **c)** *Left panel:* 3D segmentation of vascular lumen and mural cell coverage of 1^st^-8^th^-order capillaries. Note that 3 astrocytes have been mistakenly segmented as vascular lumen. *Right panel:* The mural cell coverage transitions from ensheathing pericytes to thin-strand pericytes as the capillary diameter decreases. N.B. segmentation of thin-strand pericytes is fused with endothelial cells. **d)** *Left panel:* 3D segmentation showing the vascular lumen with a yellow dashed box indicating a white matter arteriole. *Right panel:* Vascular lumen and mural cells around the arteriole and precapillary sphincter. **e)** The borders between ECs can be followed as an indentation of the vascular lumen as the tight junctions are associated with a small EC protrusion into the lumen (*).

**Supplementary Figure 2**

**a)** *Left and central panel:* Ultrastructure of a pial arteriole with elastin (white arrowheads) and collagen (black arrowheads). *Right panel:* Ultrastructure of glia limitans with collagen fibers (black arrowheads). **b)** *Left panel:* Ultrastructure of the wall of a PA shows a dense layer of elastin with a fibrillin sheath. *Right panel:* Ultrastructure of a precapillary sphincter shows where the dense elastin stops and becomes more diffuse in the fibrillin sheath. **c)** Collagen fibers can be found around vasculature, and sometimes the fibers are so dense that they show up on 3D segmentations (green). **d)** *Left panel:* Collagen fibrils (black arrowheads) are usually seen in pockets between the fibroblast and mural cells (MC). *Right panel:* Collagen fibrils can be found leading into fibroblast invaginations (fibropositors), which is putatively where they are synthesized. **e)** In the Bonney et al. 2022 study, a macrophage was mistaken for a fibroblast; however, a fibroblast exists nearby. Furthermore, this branchpoint has two mural cells encircling it, one situated at the branchpoint, and one situated at the PA (mural cells and endothelial nucleus: tinyurl.com/5953pewe), which was interpreted as a precapillary sphincter. However, that branchpoint lumen diameter is ∼8 µm *ex vivo* and it has hardly any indentation (vessel lumen: tinyurl.com/4hvkhb67), which means it is in the greyzone between a capillary and an arteriole(2). We defined precapillary sphincters as a lumen diameter of the branchpoint less than 0.8 times the lumen of the 1^st^-order capillary(7). It should be noted that this definition was set for *in vivo* measurements, and it is unclear whether all indentations are retained with the loss of blood pressure, where the microvascular-lumen diameter is instantly reduced by about 20%(7). However, about 20 µm deeper in the cortex on the same PA there is a branchpoint with clear indentation, mural cells encircling it, and followed by an exposed endothelial nucleus, indicating a precapillary sphincter and bulb (vessel lumen: tinyurl.com/ytjxtas3 and mural cells and endothelial nucleus: tinyurl.com/y76vc5vu, note that the segmentation of the green mural cell is fused with a fibroblast nucleus). **f)** Macrophages and fibroblast somas near precapillary sphincters are associated with an indentation into the PA lumen. *Left panel:* ultrastructure with macrophage (blue) and fibroblast segments indicated. *Central panel:* 3D segmentation of cells and vascular lumen. *Right panel:* 3D segmentation of vascular lumen showing indentation. **g)** Ultrastructure of the vascular wall of a pial arteriole. In places where there is no mural cell coverage the fibroblast processes come in direct contact (black arrowhead) with the endothelial-cell basement membrane.

**Supplementary Figure 3**

**a)** Large pial arterioles do not have any perivascular nerves. **b)** At the precapillary sphincter, the neuronal processes closest to the astrocyte endfeet are almost exclusively axons. Dendrites are further from away. *Left panel:* Ultrastructure and 3D segmentation of axons near the precapillary sphincter. *Right panel:* Ultrastructure and 3D segmentation of dendrites near the precapillary sphincter. **c)** The processes of many different cell types can be found close to the astrocyte endfeet, here are some examples.

**Supplementary Figure 4**

**a)** Examples of cell bodies of astrocytes, microglia, neurons, and oligodendrocyte precursor cells taking up space on one side of the vasculature, opposite to the perpendicular arranged neuronal processes and thereby polarizing the arrangement. **b)** Contractile elements (yellow arrowheads) in the mural cell encircling a 1^st^-order capillary branchpoint in the human dataset. **c)** Oligodendrocytes polarizing the perpendicular arrangement of a 1^st^-order capillary in the human dataset. **d)** Example of 4 views of the closest neuronal processes to the astrocyte endfeet of an ascending venule in the human dataset. *Left panels:* 3D segmentation of the neuronal processes, the vessel is oriented horizontally. *Central panels:* Color coding of angles of neuronal processes to the vessel orientation. *Right panels:* Quantification of the pixel distribution of angels in the 4 views.

