## Supplementary Figures and Table for "Ultrastructure of Precapillary Sphincters and the Neurovascular Unit"

Søren Grubb

**This PDF file includes:**

Figures S1 to S4

Legends for Movies S1 to S7

Supplementary Table 1

SI References

**Other supporting materials for this manuscript include the following:**

Movies S1 to S7

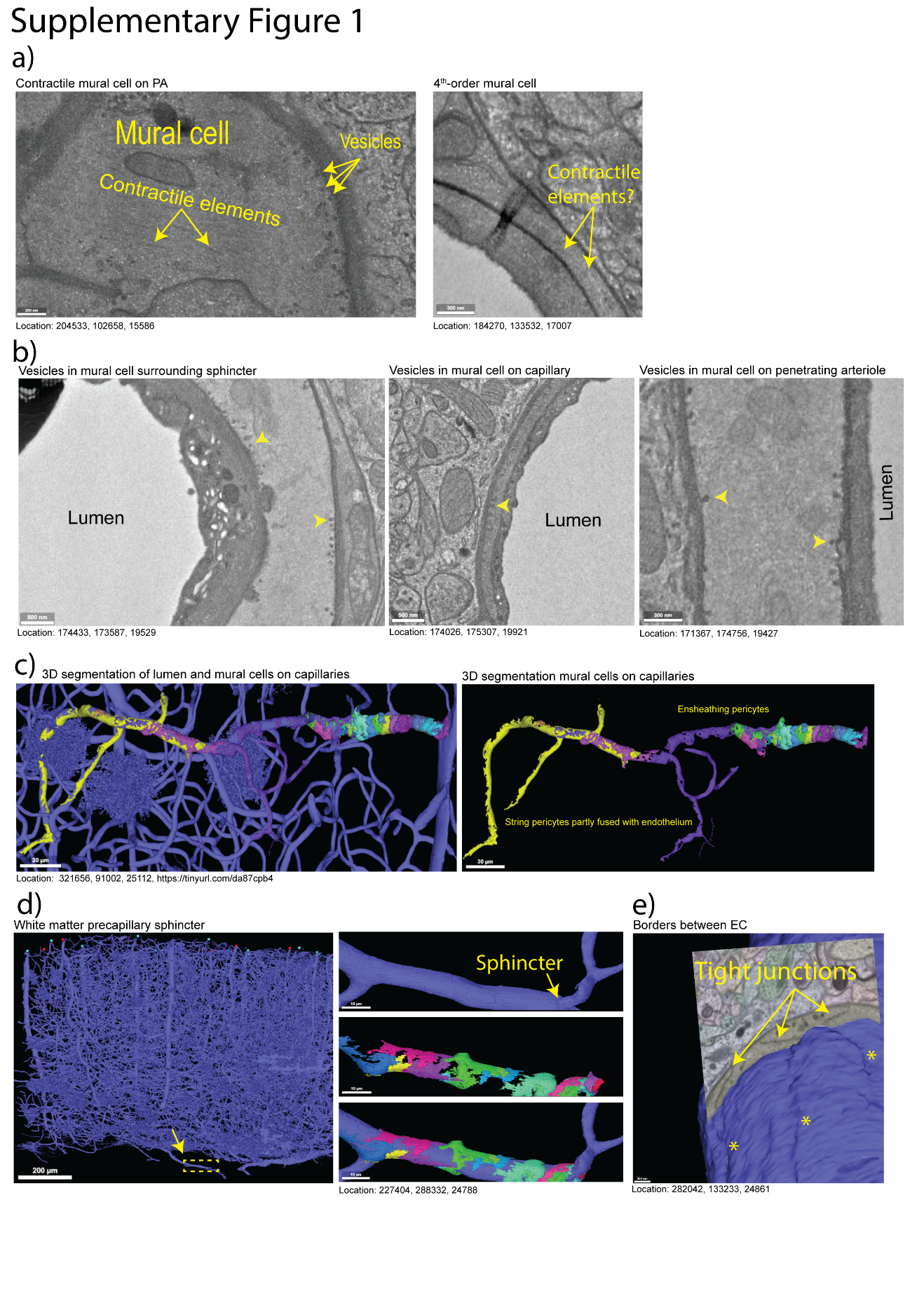

**Fig. S1.** **a)** *Left panel:* Ultrastructure of an arteriolar contractile mural cell showing contractile elements and abluminal coated vesicles. *Right panel:* Mural cell on a 4^th^-order capillary showing a few lines that could be contractile elements. **b)** *Left and right panels:* Vesicles and caveolae can be found near the luminal and abluminal membrane of mural cells on precapillary sphincters and PAs. *Central panel:* In capillaries vesicles and caveolae are mainly found on the abluminal membrane. **c)** *Left panel:* 3D segmentation of vascular lumen and mural cell coverage of 1^st^-8^th^-order capillaries. Note that 3 astrocytes have been mistakenly segmented as vascular lumen. *Right panel:* The mural cell coverage transitions from ensheathing pericytes to string pericytes as the capillary diameter decreases. N.B. segmentation of string pericytes is fused with endothelial cells. **d)** *Left panel:* 3D segmentation showing the vascular lumen with a yellow dashed box indicating a white matter arteriole. *Right panel:* Vascular lumen and mural cells around the arteriole and precapillary sphincter. **e)** The borders between ECs can be followed as an indentation of the vascular lumen as the tight junctions are associated with a small EC protrusion into the lumen (*).

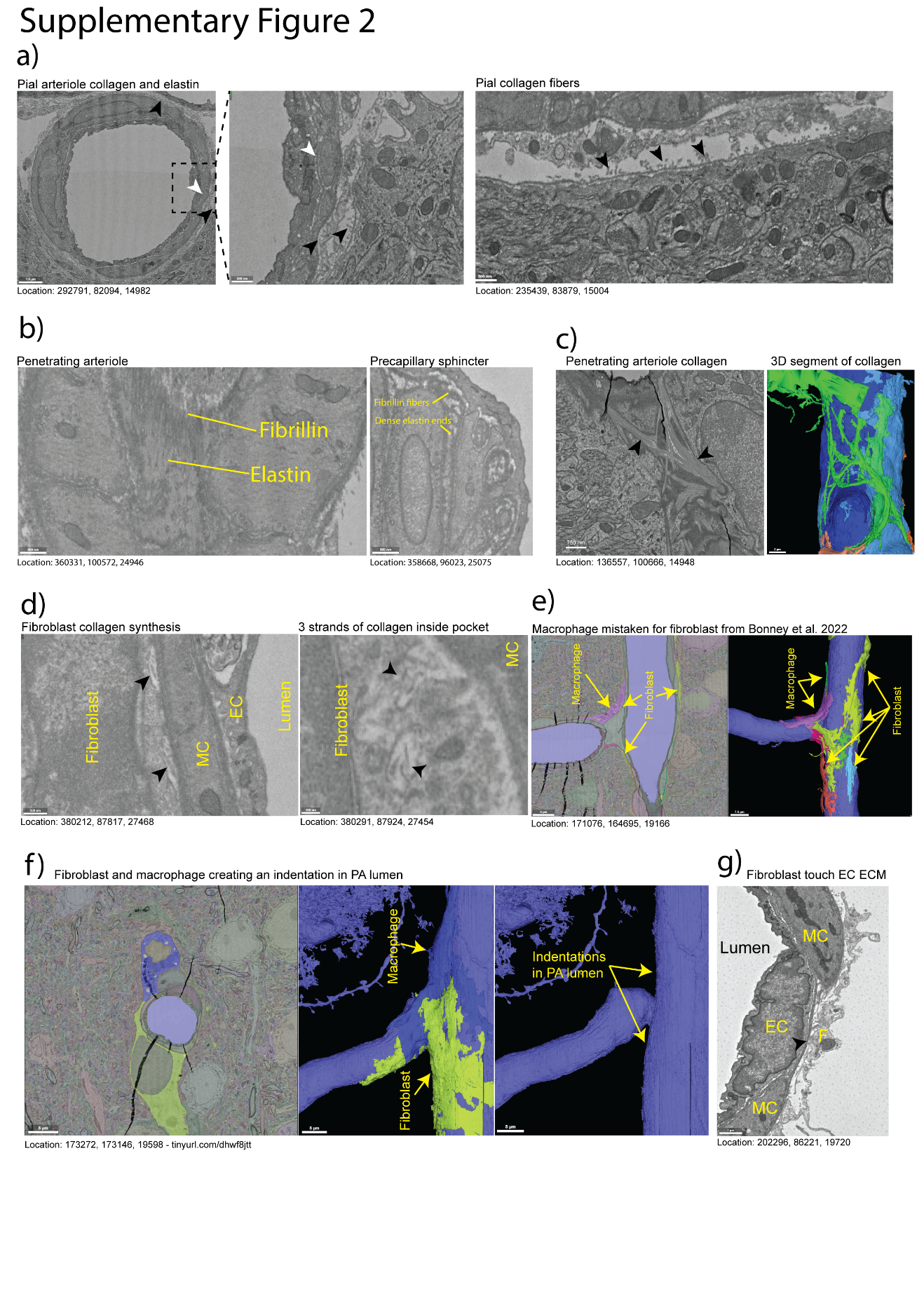

**Fig. S2.** **a)** *Left and central panel:* Ultrastructure of a pial arteriole with elastin (white arrowheads) and collagen (black arrowheads). *Right panel:* Ultrastructure of glia limitans with collagen fiber bundles (black arrowheads). **b)** *Left panel:* Ultrastructure of the wall of a PA shows a dense layer of elastin with a fibrillin sheath. *Right panel:* Ultrastructure of a precapillary sphincter shows where the dense elastin stops and becomes more diffuse in the fibrillin sheath. **c)** Collagen fibers can be found around vasculature, and sometimes the fiber bundles are so dense that they show up on 3D segmentations (green). **d)** *Left panel:* Collagen fibers (black arrowheads) are usually seen in pockets between the fibroblast and mural cells (MC). *Right panel:* Collagen fibers can be found leading into fibroblast invaginations, which is putatively where they are synthesized. **e)** In the Bonney et al. 2022 study, a macrophage was mistaken for a fibroblast; however, a fibroblast exists nearby. Furthermore, this branchpoint has two mural cells encircling it, one situated at the branchpoint, and one situated at the PA (mural cells and endothelial nucleus: [tinyurl.com/5953pewe](https://tinyurl.com/5953pewe)), which was interpreted as a precapillary sphincter. However, that branchpoint lumen diameter is ~8 µm *ex vivo* and it has hardly any indentation (vessel lumen: [tinyurl.com/4hvkhb67](https://tinyurl.com/4hvkhb67)), which means it is in the greyzone between a capillary and an arteriole(1). We defined precapillary sphincters as a lumen diameter of the branchpoint less than 0.8 times the lumen of the 1^st^-order capillary(2). It should be noted that this definition was set for *in vivo* measurements, and it is unclear whether all indentations are retained with the loss of blood pressure, where the microvascular-lumen diameter is instantly reduced by about 20%(2). However, about 20 µm deeper in the cortex on the same PA there is a branchpoint with clear indentation, mural cells encircling it, and followed by an exposed endothelial nucleus, indicating a precapillary sphincter and bulb (vessel lumen: [tinyurl.com/ytjxtas3](http://tinyurl.com/ytjxtas3) and mural cells and endothelial nucleus: [tinyurl.com/y76vc5vu](https://tinyurl.com/y76vc5vu), note that the segmentation of the green mural cell is fused with a fibroblast nucleus). **f)** Macrophages and fibroblast somas near precapillary sphincters are associated with an indentation into the PA lumen. *Left panel:* ultrastructure with macrophage (blue) and fibroblast segments indicated. *Central panel:* 3D segmentation of cells and vascular lumen. *Right panel:* 3D segmentation of vascular lumen showing indentation. **g)** Ultrastructure of the vascular wall of a pial arteriole. In places where there is no mural cell coverage the fibroblast processes come in direct contact (black arrowhead) with the endothelial-cell basement membrane.

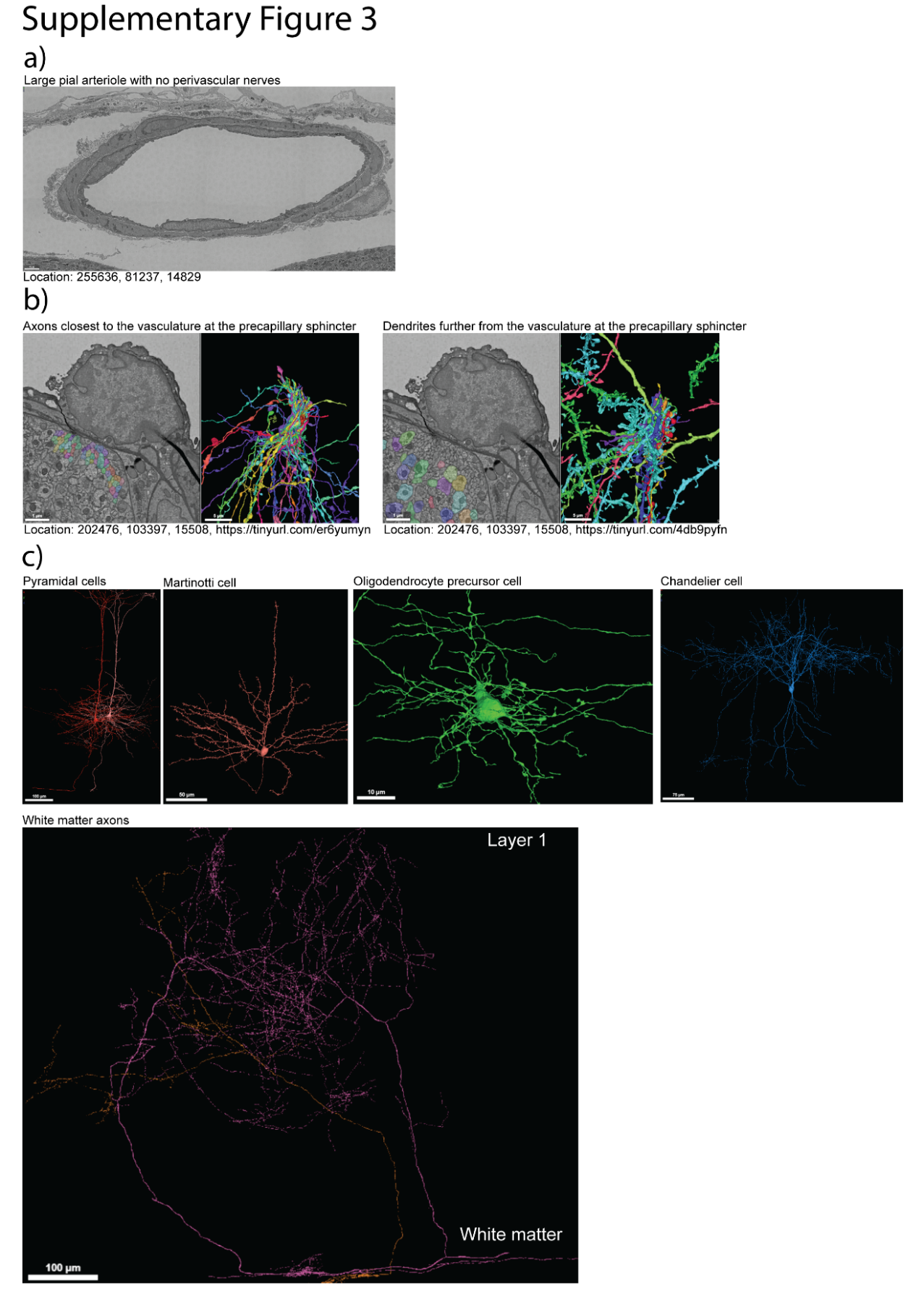

**Fig. S3**. **a)** Large pial arterioles do not have any perivascular nerves. **b)** At the precapillary sphincter, the neuronal processes closest to the astrocyte endfeet are almost exclusively axons. Dendrites are further from away. *Left panel:* Ultrastructure and 3D segmentation of axons near the precapillary sphincter. *Right panel:* Ultrastructure and 3D segmentation of dendrites near the precapillary sphincter. **c)** The processes of many different cell types can be found close to the astrocyte endfeet, here are some examples.

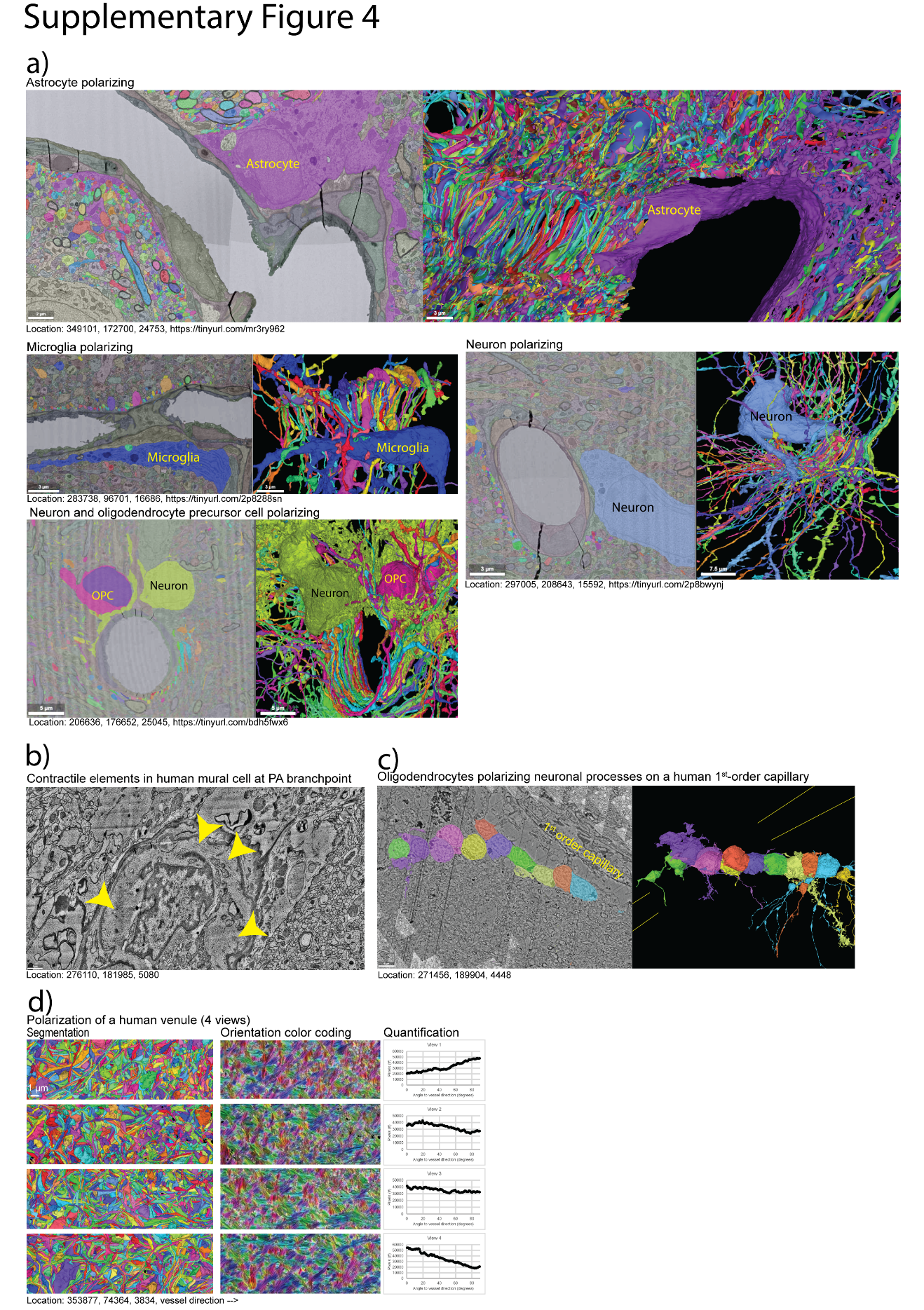

**Fig. S4.** **a)** Examples of cell bodies of astrocytes, microglia, neurons, and oligodendrocyte precursor cells taking up space on one side of the vasculature, opposite to the perpendicular arranged neuronal processes and thereby polarizing the arrangement. **b)** Contractile elements (yellow arrowheads) in the mural cell encircling a 1^st^-order capillary branchpoint in the human dataset. **c)** Oligodendrocytes polarizing the perpendicular arrangement of a 1^st^-order capillary in the human dataset. **d)** Example of 4 views of the closest neuronal processes to the astrocyte endfeet of an ascending venule in the human dataset. *Left panels:* 3D segmentation of the neuronal processes, the vessel is oriented horizontally. *Central panels:* Color coding of angles of neuronal processes to the vessel orientation. *Right panels:* Quantification of the pixel distribution of angels in the 4 views.

Movie S1 (separate file). Showing the morphology of mural cells (various colors) encircling a PA, precapillary sphincter and 1^st^ - 3^rd^-order capillaries. The vascular lumen is light blue and endothelial cells are not shown. Notice how the morphology transitions from mural cells with a more flat soma and 3-4 processes encircling the PA (red, purple and cyan), while the ensheathing pericyte at the precapillary sphincter (cyan) and the ensheathing pericyte covering the 1^st^- and 2^nd^-order capillaries (pink) have more protruding somas and several processes ensheathing the capillaries. The thin-strand pericyte on the 3^rd^-order capillary also have a protruding soma, but the processes are increasinglu oriented along the capillary rather than encircling it. The location in the MICrONS dataset is: 225010, 158639, 19156. N.B. the cyan segmentation contains both an ensheathing pericyte at the precapillary sphincter, parts of a PA mural cell and parts of a neuronal process.

Movie S2 (separate file). Showing individual layers in the MICrONS dataset serial EM of the PA and precapillary sphincter shown in Figure 1c. The video is paused at different layers to show the presence of collagen fibers at the bulb (part of 1^st^-order capillary), the location of endothelial cells, mural cells, fibroblasts, a macrophage and astrocytes, contractile elements, differences in glycocalyx density, pinocytic vesicles, elastin, myoendothelial junctions and junctions between mural cells (myo-myo junctions). The location in the MICrONS dataset is: 203779, 103491, 15475.

Movie S3 (separate file). Showing annotations (yellow dots) of myoendothelial junctions and/or peg-and-socket junctions at a PA, a precapillary sphincter, a 1^st^ order capillary, 2^nd^- and 3^rd^-order capillaries. Mural cells (various colors) and vascular lumen (light blue) have a low opacity to make the annotations visible. Mural cell somas are marked by yellow asterisks. Notice how relative to the PA and 1^st^-order capillary the precapillary sphincter has many myoendothelial junctions. This is also depicted in Figure 1d and the location in the MICrONS dataset is: 349084, 92330, 25337.

Movie S4 (separate file). Showing individual layers in the MICrONS dataset serial EM of a vascular wall of a PA where endothelial cells, a mural cells, a fibroblast, the PVS and astrocytic endfeet are indicated. The video pauses at a location where pockets in the fibroblasts contains leading end of collagen strands (fibrils); however, this can be appreciated best by playing and rewinding the movie while following the collagen fibers. Collagen fibers can also be seen traversing the PVS. This is also depicted in Supp. Fig. 2d and the location in the MICrONS dataset is: 380203, 87715, 27397.

Movie S5 (separate file). Showing individual layers in the MICrONS dataset serial EM while moving to follow the astrocyte endfeet around a PA with PVS. The astrocyte segmentations are in various colors, but the green astrocyte is mistakenly segmented together with mural cells. Not all astrocyte segmentations are shown. Notice how the astrocyte somas form a part of the endfeet and how the astrocytic endfeet overlap substantially. This is also depicted in Figure 2a and the location in the MICrONS dataset is: 201978, 82707, 24442.

Movie S6 (separate file). Showing 3D segmentations of mouse neuronal processes (various colors) on the abluminal side of astrocytic endfeet for a shallow PA and a shallow 1^st^ order capillary with PVS, a deep PA without PVS, a shallow venule without PVS and a deep PA with PVS. The vasculature and PVS are not shown, but the vascular direction is indicated (horizontally). Each movie starts with cross-sectional view, increases the focal depth, turns to a side-view and tilts.

Movie S7 (separate file). Showing 3D segmentations of human neuronal processes (various colors) on the abluminal side of astrocytic endfeet for a 1^st^ order capillary (Location in the H01 dataset: 277102, 180552, 5259) and a venule (Location in the H01 dataset: 353116, 70815, 3916). The vasculature is not shown, but the vascular direction is indicated (horizontally). Each movie starts with cross-sectional view, increases the focal depth, turns to a side-view and tilts.

### Supplementary Table 1 – quantification of precapillary sphincters:

| Arteriole# | Branch# | Coordinates | Sphincter Ø | 1. Order Ø | Ratio | Classification |
| --- | --- | --- | --- | --- | --- | --- |
| 1 | 1 | 74595, 108426, 19742 | 3.58 | 7.14 | 0.50 | Sphincter |
|  | 2 | 75268, 117870, 19100 | 4.98 | 6.52 | 0.76 | Sphincter |
|  | 3 | 72732, 153978, 19551 | 4.84 | 6.25 | 0.77 | Sphincter |
| 2 | 1 | 170006, 165066, 19289 | 7.72 | 7.98 | 0.97 | Not Sphincter |
|  | 2 | 173800, 173543, 19554 | 4.67 | 5.94 | 0.79 | Sphincter |
|  | 3 | 173787, 212162, 18394 | NA | NA | NA | Arteriole |
|  | 4 | 173755, 209460, 17937 | NA | NA | NA | Arteriole |
|  | 5 | 172137, 206164, 17850 | 2.97 | 3.28 | 0.91 | Not Sphincter |
|  | 6 | 168645, 199074, 17966 | 5.80 | 6.80 | 0.85 | Not Sphincter |
| 3 | 1 | 113960, 92549, 23418 | 2.39 | 4.37 | 0.55 | Sphincter |
|  | 2 | 122150, 181802, 22428 | NA | NA | NA | Arteriole |
|  | 3 | 121444, 184487, 22724 | 7.91 | 7.39 | 1.07 | Not Sphincter |
|  | 4 | 121747, 195362, 23039 | 3.18 | 3.92 | 0.81 | Not Sphincter |
| 4 | 1 | 210284, 109979, 25409 | NA | NA | NA | Bifurcation |
|  | 2 | 204773, 163891, 24599 | 6.04 | 6.36 | 0.95 | Not Sphincter |
|  | 3 | 202486, 172266, 24849 | 4.93 | 5.72 | 0.86 | Not Sphincter |
|  | 4 | 204976, 179241, 24959 | 8.86 | 8.64 | 1.02 | Not Sphincter |
|  | 5 | 212398, 190774, 24360 | 4.70 | 6.10 | 0.77 | Sphincter |
|  | 6 | 210644, 204189, 24922 | 6.16 | 6.06 | 1.02 | Not Sphincter |
|  | 7 | 206126, 207593, 24682 | 7.36 | 7.04 | 1.05 | Not Sphincter |
|  | 8 | 196045, 207297, 24256 | 2.15 | 2.54 | 0.85 | Not Sphincter |
| 5 | 1 | 357846, 94680, 25093 | 4.95 | 7.24 | 0.68 | Sphincter |
|  | 2 | 357048, 106431, 25451 | 4.77 | 5.65 | 0.84 | Not Sphincter |
|  | 3 | 359213, 106537, 24991 | 6.81 | 8.93 | 0.76 | Sphincter |
|  | 4 | 351485, 169041, 25054 | 3.76 | 4.11 | 0.92 | Not Sphincter |
|  | 5 | 348820, 173983, 24917 | 5.22 | 5.65 | 0.92 | Not Sphincter |
|  | 6 | 339946, 194450, 25441 | 4.71 | 4.84 | 0.97 | Not Sphincter |
|  | 7 | 340570, 204125, 25050 | NA | NA | NA | Bifurcation |
|  | 8 | 335837, 212486, 25656 | 3.26 | 3.67 | 0.89 | Not Sphincter |
|  | 9 | 333938, 215257, 25680 | 3.59 | 4.04 | 0.89 | Not Sphincter |
|  | 10 | 331871, 227197, 25736 | 6.72 | 6.34 | 1.06 | Not Sphincter |
| 7 | 1 | 225627, 153955, 19298 | 5.80 | 7.98 | 0.73 | Sphincter |
|  | 2 | 231199, 199952, 20491 | 6.14 | 6.88 | 0.89 | Not Sphincter |
|  | 3 | 236667, 207676, 19947 | 8.24 | 8.32 | 0.99 | Not Sphincter |
|  | 4 | 234677, 213898, 20550 | NA | NA | NA | Arteriole |
|  | 5 | 233765, 214506, 20427 | 11.18 | 8.48 | 1.32 | Not Sphincter |
| 8 | 1 | 286906, 97518, 16687 | 3.36 | 3.12 | 1.08 | Not Sphincter |
|  | 2 | 283959, 126535, 17497 | NA | NA | NA | Arteriole |
|  | 3 | 286643, 131713, 17327 | NA | NA | NA | Arteriole |
|  | 4 | 284275, 141142, 17436 | 3.55 | 3.20 | 1.11 | Not Sphincter |
| 9 | 1 | 236463, 129244, 16871 | 6.62 | 5.62 | 1.18 | Not Sphincter |
|  | 2 | 241777, 132659, 17301 | NA | NA | NA | Arteriole |
|  | 3 | 246791, 154761, 18087 | 3.47 | 4.17 | 0.83 | Not Sphincter |
|  | 4 | 254674, 158598, 18859 | 4.45 | 4.45 | 1.00 | Not Sphincter |
|  | 5 | 256219, 158816, 19143 | 4.61 | 4.91 | 0.94 | Not Sphincter |
| 10 | 1 | 203857, 103574, 15476 | 3.30 | 4.22 | 0.78 | Sphincter |
|  | 2 | 204554, 134469, 15410 | 5.81 | 4.34 | 1.34 | Not Sphincter |
|  | 3 | 200877, 137255, 15208 | NA | NA | NA | Bifurcation |
| 11 | 1 | 316214, 87729, 16498 | 3.12 | 3.61 | 0.87 | Not Sphincter |
|  | 2 | 316857, 100964, 17635 | 4.65 | 6.35 | 0.73 | Sphincter |
|  | 3 | 314010, 112441, 17739 | 4.23 | 5.45 | 0.78 | Sphincter |
|  | 4 | 317051, 149938, 18634 | 3.90 | 3.92 | 0.99 | Not Sphincter |
|  | 5 | 311308, 158316, 18465 | NA | NA | NA | Arteriole |
| 12 | 1 | 153011, 97361, 23487 | 3.89 | 4.19 | 0.93 | Not Sphincter |
|  | 2 | 154632, 96935, 23378 | 3.48 | 4.64 | 0.75 | Sphincter |
|  | 3 | 156408, 155386, 23591 | NA | NA | NA | Arteriole |
|  | 4 | 157740, 160353, 23861 | 4.06 | 4.43 | 0.92 | Not Sphincter |
|  | 5 | 159002, 167789, 23673 | 4.79 | 5.05 | 0.95 | Not Sphincter |
|  | 6 | 160038, 176438, 23683 | NA | NA | NA | Bifurcation |
| 13 | 1 | 311988, 189957, 14564 | NA | NA | NA | Arteriole |
|  | 2 | 307513, 190698, 13989 | NA | NA | NA | Bifurcating |
|  | 3 | 297372, 200369, 15150 | 4.42 | 6.77 | 0.65 | Sphincter |
|  | 4 | 293972, 207619, 15405 | 3.75 | 5.75 | 0.65 | Sphincter |
|  | 5 | 285692, 214274, 14962 | 5.42 | 6.73 | 0.80 | Not Sphincter |
| 14 | 1 | 324970, 136938, 11893 | NA | NA | NA | Arteriole |
|  | 2 | 326803, 147409, 12565 | 2.86 | 4.99 | 0.57 | Sphincter |
|  | 3 | 336099, 168261, 12430 | 5.11 | 7.02 | 0.73 | Sphincter |
| 15 | 1 | 232832, 101719, 8418 | 3.57 | 4.66 | 0.77 | Sphincter |
|  | 2 | 229982, 132612, 8198 | 4.67 | 5.02 | 0.93 | Not Sphincter |
|  | 3 | 230329, 139604, 8176 | 4.43 | 4.55 | 0.97 | Not Sphincter |
|  | 4 | 229868, 147087, 8382 | 3.29 | 2.95 | 1.11 | Not Sphincter |
| 16 | 1 | 117487, 101356, 8596 | 2.65 | 3.25 | 0.82 | Not Sphincter |
|  | 2 | 121799, 110848, 8747 | 3.14 | 3.21 | 0.98 | Not Sphincter |
|  | 3 | 120183, 115464, 8974 | NA | NA | NA | Arteriole |
| 17 | 1 | 91516, 146482, 10923 | NA | NA | NA | Arteriole |
|  | 2 | 99013, 187604, 10661 | 6.44 | 6.55 | 0.98 | Not Sphincter |
|  | 3 | 101478, 198669, 11548 | 4.15 | 5.60 | 0.74 | Sphincter |
|  | 4 | 108500, 208824, 11448 | 5.94 | 5.16 | 1.15 | Not Sphincter |
|  | 5 | 114477, 217696, 11970 | 4.68 | 4.89 | 0.96 | Not Sphincter |
|  | 6 | 114477, 217696, 11970 | 4.66 | 5.03 | 0.92 | Not Sphincter |
| 18 | 1 | 325187, 137311, 11863 | NA | NA | NA | Arteriole |
|  | 2 | 326316, 146821, 12542 | 2.89 | 4.63 | 0.62 | Sphincter |
|  | 3 | 335957, 169954, 11837 | 5.41 | 6.31 | 0.86 | Not Sphincter |
| 19 | 1 | 229089, 91841, 13522 | NA | NA | NA | Arteriole |
|  | 2 | 230532, 133200, 13774 | 5.04 | 4.84 | 1.04 | Not Sphincter |
|  | 2 | 235692, 93125, 12861 | 5.35 | 4.74 | 1.13 | Not Sphincter |
|  | 3 | 241644, 91523, 12894 | 1.53 | 3.58 | 0.43 | Sphincter |
|  | 3 | 229749, 144984, 13996 | 4.80 | 5.53 | 0.87 | Not Sphincter |
|  | 4 | 232541, 159579, 14209 | NA | NA | NA | Bifurcation |
|  | 5 | 229886, 167626, 14318 | 6.33 | 6.01 | 1.05 | Not Sphincter |
|  | 6 | 232026, 181203, 14557 | 4.74 | 5.04 | 0.94 | Not Sphincter |
| 20 | 1 | 211221, 104394, 14668 | 4.74 | 4.37 | 1.09 | Not Sphincter |
|  | 2 | 212002, 135493, 14535 | 5.06 | 5.40 | 0.94 | Not Sphincter |
| 21 | 1 | 170378, 112436, 8536 | 2.47 | 3.46 | 0.71 | Sphincter |
|  | 2 | 169601, 136370, 8999 | 4.88 | 4.89 | 1.00 | Not Sphincter |
|  | 3 | 174295, 154198, 9048 | NA | NA | NA | Arteriole |
|  | 4 | 173463, 159538, 9883 | 5.71 | 5.19 | 1.10 | Not Sphincter |
|  | 5 | 167585, 164896, 10445 | 4.71 | 4.73 | 1.00 | Not Sphincter |

Percent sphincters per arteriole branch

|  | Percent sphincters (%) |
| --- | --- |
| 1. branch | 35 |
| 2. branch | 28.5714 |
| 3. branch | 35 |
| 4. branch | 7.14286 |
| 5. branch | 9.09091 |
| >5. branch | 0 |
